# Neutralization of Omicron BA.4/BA.5 and BA.2.75 by Booster Vaccination or BA.2 Breakthrough Infection Sera

**DOI:** 10.1101/2022.08.04.502716

**Authors:** Xun Wang, Jingwen Ai, Xiangnan Li, Xiaoyu Zhao, Jing Wu, Haocheng Zhang, Xing He, Chaoyue Zhao, Rui Qiao, Minghui Li, Yuchen Cui, Yanjia Chen, Lulu Yang, Zixin Hu, Chenqi Xu, Wenhong Zhang, Pengfei Wang

**Affiliations:** State Key Laboratory of Genetic Engineering, Shanghai Institute of Infectious Disease and Biosecurity, School of Life Sciences, Fudan University, Shanghai, China; Department of Infectious Diseases, Shanghai Key Laboratory of Infectious Diseases and Biosafety Emergency Response, National Medical Center for Infectious Diseases, Huashan Hospital, Fudan University, Shanghai, China; State Key Laboratory of Genetic Engineering, Collaborative Innovation Center for Genetics and Development, School of Life Sciences and Human Phenome Institute, Zhangjiang Fudan International Innovation Center, Fudan University, Shanghai, China; Institute of Biochemistry and Cell Biology, Shanghai Institutes for Biological Sciences, Chinese Academy of Sciences, Shanghai, China; Artificial Intelligence Innovation and Incubation Institute, Fudan University, Shanghai, China; National Clinical Research Center for Aging and Medicine, Huashan Hospital, Fudan University, Shanghai, China

## Abstract

Many new Omicron sub-lineages have been reported to evade neutralizing antibody response, including BA.2, BA.2.12.1, BA.4 and BA.5. Most recently, another emerging sub-lineage BA.2.75 has been reported in multiple countries. In this study, we constructed a comprehensive panel of pseudoviruses (PsVs), including wild-type, Delta, BA.1, BA.1.1, BA.2, BA.3, BA.2.3.1, BA.2.10.1, BA.2.12.1, BA.2.13, BA.2.75 and BA.4/BA.5, with accumulate coverage reached 91% according to the proportion of sequences deposited in GISAID database since Jan 1^st^, 2022. We collected serum samples from healthy adults at day14 post homologous booster with BBIBP-CorV, or heterologous booster with ZF2001, primed with two doses of BBIBP-CorV, or from convalescents immunized with three-dose inactivated vaccines prior to infection with Omicron BA.2, and tested their neutralization activity on this panel of PsVs. Our results demonstrated that all Omicron sub-lineages showed substantial evasion of neutralizing antibodies induced by vaccination and infection, although BA.2.75 accumulated the largest number of mutations in its spike, BA.4 and BA.5 showed the strongest serum escape. However, BA.2 breakthrough infection could remarkably elevated neutralization titers against all different variants, especially titers against BA.2 and its derivative sub-lineages.

## Body text

With the continued mutation of severe acute respiratory syndrome coronavirus 2 (SARS-CoV-2) Omicron variant, many new Omicron sub-lineages have been reported to evade neutralizing antibodies induced by both vaccination and infection, including BA.2, BA.2.12.1, BA.4 and BA.5.^1-3^ Most recently, another emerging sub-lineage BA.2.75,^4^ carrying nine additional mutations in spike compared to BA.2 (Figure 1A), has been reported in multiple countries. Our previous study showed that homologous or heterologous booster can remarkably reduce Omicron BA.1, BA.1.1, BA.2 and BA.3 escape from neutralizing antibodies,^5^ but a comprehensive neutralization assessment of booster vaccination or breakthrough infection sera against all the distinct emerging Omicron sub-lineages is still lacking.

**Figure 1.**
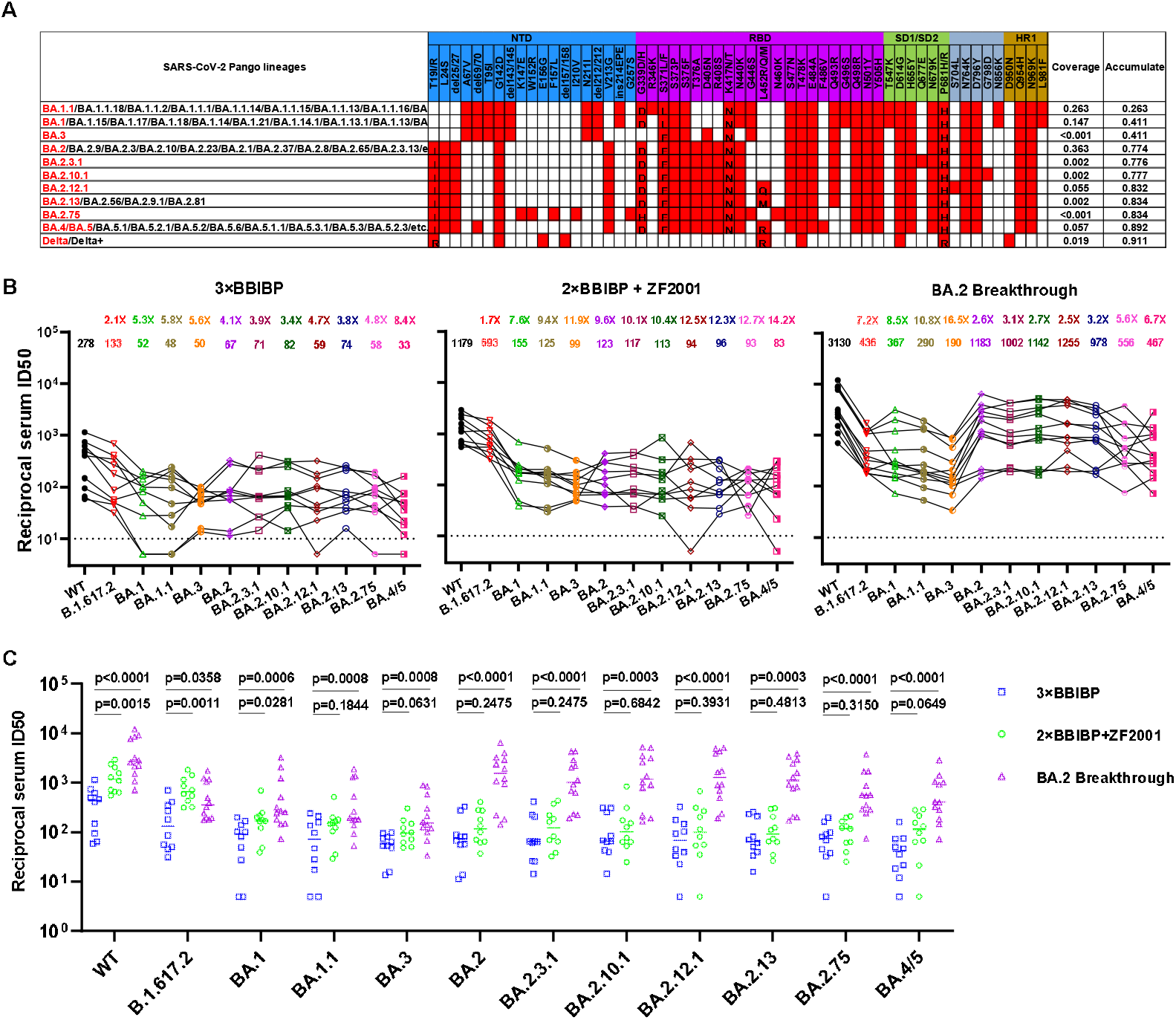
Characteristics and sera neutralization of the Omicron sub-lineages. **(A)** Prevalence and Spike mutations of the Omicron sub-lineages and Delta variant based on all the sequences available on GISAID since Jan 1^st^, 2022. **(B)** Neutralization of pseudotyped WT (D614G), Delta and Omicron sub-lineage viruses by sera collected from individuals at day 14 after vaccinated with a BBIBP-CorV homologous booster or with a ZF001 heterologous booster dose following two doses of BBIBP-CorV, or infected by BA.2 virus after three doses of BBIBP-CorV vaccination. For all panels, values above the symbols denote geometric mean titer and the fold-change was calculated by comparing the titer to WT. **(C)** In parallel comparison of neutralization titers against distinct SARS-CoV-2 variants by sera collected from individuals at day 14 after vaccinated with homologous or heterologous booster, or breakthrough infected with BA.2 virus. *P* values were determined by using Multiple Mann-Whitney tests.

Here, apart from the four Omicron sub-lineage pseudoviruses (PsVs) we already had from our previous study, we further constructed a panel of PsVs on the top of BA.2, including BA.2.3.1, BA.2.10.1, BA.2.12.1, BA.2.13, BA.2.75 and BA.4/BA.5. Some of these Omicron sub-lineages bear identical spike protein with many other sub-lineages evolved from BA.1, BA.2 or BA.5, and thus our virus panel can represent many more Omicron sub-lineages regarding their neutralization evasion levels (Figure 1A). We also included Delta (B.1.617.2) variant in this study due to its recent replacement by Omicron. Taken all these variants and their representing variants into account, their accumulate coverage reached 91% according to the proportion of sequences deposited in GISAID database since Jan 1^st^, 2022 (Figure 1A), which is the most comprehensive panel of Omicron sub-lineages tested as we know.

We collected serum samples from healthy adults at day14 post homologous booster with BBIBP-CorV, or heterologous booster with ZF2001, primed with two doses of BBIBP-CorV (Table S1), and tested their neutralization activity on this panel of PsVs. As shown in Figure 1B and Fig. S1, the homologous booster group (3×BBIBP, n=10) had a neutralizing geometric mean titer (GMT) against WT of 278, with 2.1- to 8.4-fold reduction against Delta and Omicron sub-lineages. For the heterologous booster group (2×BBIBP + ZF2001, n=10), this cohort had higher neutralizing titers with GMTs of 1179, 693, 155, 125, 99, 123, 117, 94, 96, 93 and 83 against WT, Delta, BA.1, BA.1.1, BA.3, BA.2, BA.2.3.1, BA.2.10.1, BA.2.12.1, BA.2.13, BA.2.75 and BA.4/5, respectively. Although these numbers amount to 1.7- to 14.2-fold reductions in potency for the Delta and Omicron sub-lineages compared to WT, nearly all samples retained detectable neutralizing activity against these distinct variants. Of note, although BA.2.75 accumulated the largest number of mutations in its spike, BA.4 and BA.5 showed the strongest serum escape in both the homologous and heterologous booster groups.

The SARS-CoV-2 BA.2 variant has led to an increasing number of breakthrough infections in China. To gain further insight into their chance of re-infection by new Omicron sub-lineages, we recruited 12 convalescents immunized with three-dose inactivated vaccines prior to infection with Omicron BA.2 and evaluated their serum samples at day 14 post-infection on the same panel of PsVs. We found that BA.2 breakthrough infection significantly increased neutralizing antibody to higher titers with GMTs of 3130, 436, 367, 290, 190, 1183, 1002, 1142, 1255, 978, 556 and 467 against WT, B.1.617.2, BA.1, BA.1.1, BA.3, BA.2, BA.2.3.1, BA.2.10.1, BA.2.12.1, BA.2.13, BA.2.75 and BA.4/5 (Figure 1B). For BA.2, its derivative variants and BA.4/5, the reduction levels compared to WT in the breakthrough infection group were lower than those of the homologous and heterologous vaccine booster groups. While for Delta, BA.1, BA.1.1 and BA.3, the reduction levels were much higher, which may be associated with the antigenic difference between Omicron BA.2 and these other variants.

To further understand the differences between vaccination and BA.2 breakthrough infection, we compared in parallel the serum neutralization titers of homologous booster, heterologous booster, and BA.2 breakthrough infection, against different viruses. Heterologous booster exhibited higher titers than homologous booster against WT and Delta variant. However, the neutralization titers for Omicron sub-lineages showed no difference between homologous and heterologous (RBD-subunit) boosters, which could be attributed to the large number of mutations accumulated in Omicron variants, especially those in RBD. More importantly, we found that BA.2 breakthrough infection significantly increased neutralizing antibody titers to relatively high levels compared with homologous and heterologous booster vaccination against almost all variants (Figure 1C). For the BA.2 breakthrough infection sera, we further compared the neutralization titers against BA.2 with those of the other variants. BA.2.75 and BA.4/5, with several additional mutations on the top of BA.2, showed significant lower titers than BA.2. While for the other sub-lineages like BA.2.3.1, BA.2.10.1 and BA.2.12.1, which are derived from BA.2 with only one additional mutation, we observed similar response to the breakthrough infection sera compared to BA.2 (Fig. S2).

Taken together, our results demonstrated that all Omicron sub-lineages showed substantial evasion of neutralizing antibodies induced by vaccination, with BA.4/5 to be the most significant one. However, BA.2 breakthrough infection could remarkably elevated neutralization titers against all different variants, especially titers against BA.2 and its derivative sub-lineages.

## Supplementary Methods

### Serum samples

Sera from individuals who received three doses of BBIBP-CorV or two doses of BBIBP-CorV plus ZF2001 vaccine were collected at Huashan Hospital, Fudan University 14 days after the final dose. Sera were also obtained from patients after 14 days of SARS-CoV-2 breakthrough infection caused by Omicron BA.2 variant after immunizing with three-dose inactivated vaccines (CoronaVac or BBIBP-CorV). All collections were conducted according to the guidelines of the Declaration of Helsinki and approved by the Institutional Review Board of the Ethics Committee of Huashan Hospital (2021-041 and 2021-749). All the participants provided written informed consents.

### Construction and production of variant pseudoviruses

Plasmids encoding the WT (D614G) SARS-CoV-2 spike and Omicron sub-lineage spikes, as well as the spikes with single or combined mutations were synthesized. Expi293F cells were grown to 3×10^6^/mL before transfection with the indicated spike gene using Polyethylenimine (Polyscience). Cells were cultured overnight at 37 °C with 8% CO_2_ and VSV-G pseudo-typed ΔG-luciferase (G*ΔG-luciferase, Kerafast) was used to infect the cells in DMEM at a multiplicity of infection of 5 for 4 h before washing the cells with 1×DPBS three times. The next day, the transfection supernatant was collected and clarified by centrifugation at 300g for 10 min. Each viral stock was then incubated with 20% I1 hybridoma (anti-VSV-G; ATCC, CRL-2700) supernatant for 1 h at 37 °C to neutralize the contaminating VSV-G pseudotyped ΔG-luciferase virus before measuring titers and making aliquots to be stored at -80 °C.

### Pseudovirus neutralization assays

Neutralization assays were performed by incubating pseudoviruses with serial dilutions of monoclonal antibodies or sera, and scored by the reduction in luciferase gene expression. In brief, Vero E6 cells were seeded in a 96-well plate at a concentration of 2×10^4^ cells per well. Pseudoviruses were incubated the next day with serial dilutions of the test samples in triplicate for 30 min at 37 °C. The mixture was added to cultured cells and incubated for an additional 24 h. The luminescence was measured by Luciferase Assay System (Beyotime). IC_50_ was defined as the dilution at which the relative light units were reduced by 50% compared with the virus control wells (virus + cells) after subtraction of the background in the control groups with cells only. The IC_50_ values were calculated using nonlinear regression in GraphPad Prism.

### Analysis of spike mutations and prevalence of SARS-CoV-2 variants

The coverage of each lineage group was summarized by counting the submission records of each lineage in the same group since 2022 from GISAID database (Accessed on 30 July, 2022). The lineages were grouped together if they share same mutations in at least 60% submitted sequences in spike protein. Lineages with prefix “AY” were grouped as Delta+ lineage. The mutation data of each lineage were retrieved using R package outbreakinfo with lookupSublineages() and getMutationsByLineage() functions. We then filtered and counted the submission records of each lineage and calculated the coverage of each lineage group. Only lineages with both mutation data and submission records were taken into account.

---

**Table S1.** Baseline characteristics of enrolled participants, including Omicron BA.2 breakthrough infection, BBIBP-CorV homologous booster group and BBIBP-CorV/ ZF2001 heterologous booster group.

|  | Omicron BA.2<br>breakthrough<br>infection<br>(n=12) | BBIBP-CorV<br>homologous<br>booster dose<br>(n=10) | BBIBP-CorV/ZF2001<br>heterologous booster<br>dose (n=10) | P value |
| --- | --- | --- | --- | --- |
| Age(years),<br>median(range) | 26(21-31) | 28(19-33) | 22.5(23-51) | 0.02 |
| Male, n(%) | 4(33.33%) | 4(40.00%) | 3(30.00%) | 0.90 |
| BMI(kg/m<sup>2</sup>),mean(SD) | 22.02(2.52) | 20.87(2.38) | 21.18(2.4) | 0.56 |
| Comorbidities(%) |  |  |  |  |
| Any, n(%) | 1(8.30%) | 1(10.00%) | 0(0.00%) | 0.63 |
| Tumor, n(%) | 0(0.00%) | 1(10.00%) | 0(0.00%) |  |
| Others, n(%) | 1(8.30%) | 0(0.00%) | 0(0.00%) | 0.63 |

**Figure S1.**
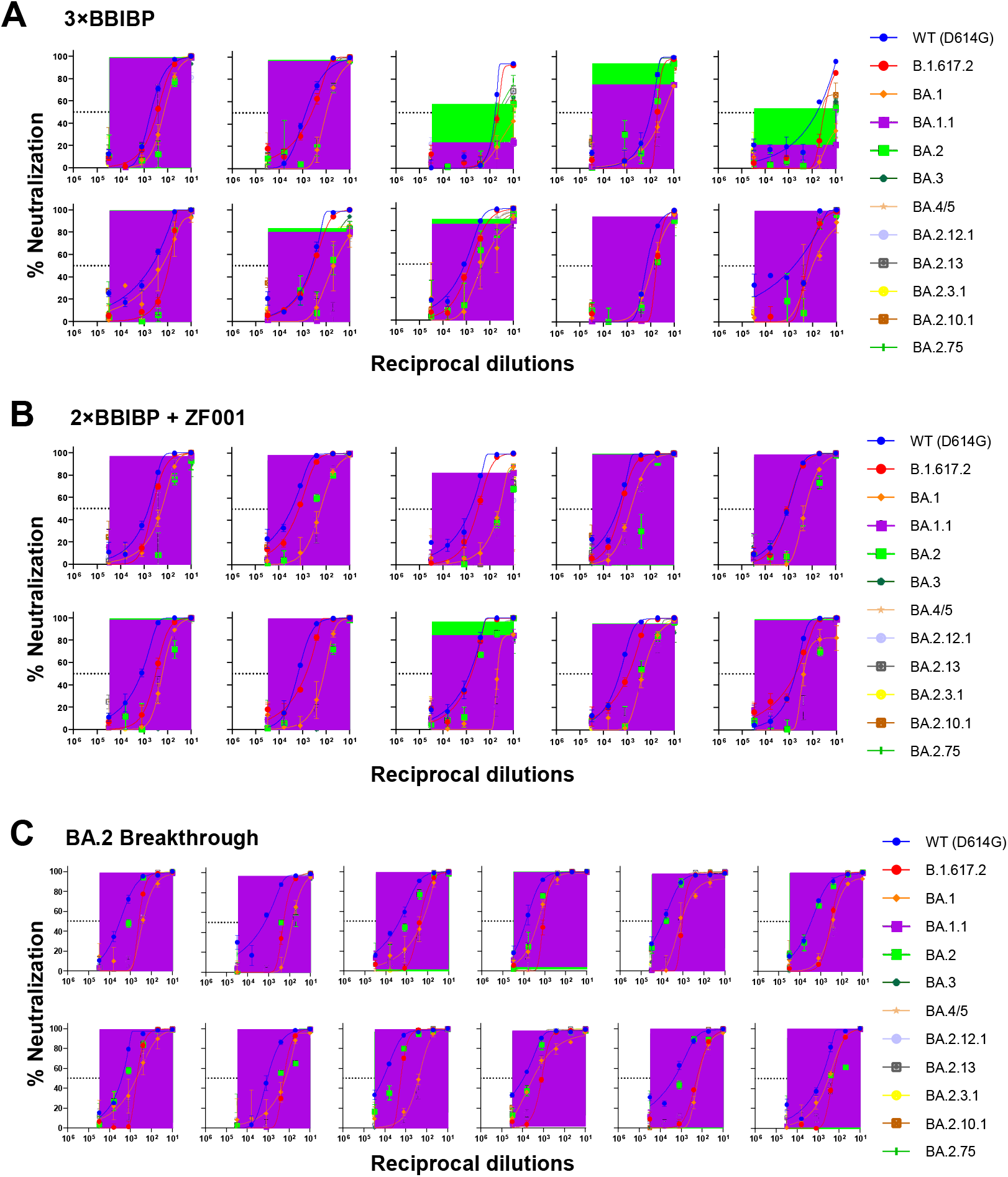
Neutralization curves for sera collected at day 14 post the BBIBP-CorV (A) or ZF001 booster dose (B) or BA.2 breakthrough infection (C)

**Figure S2.**
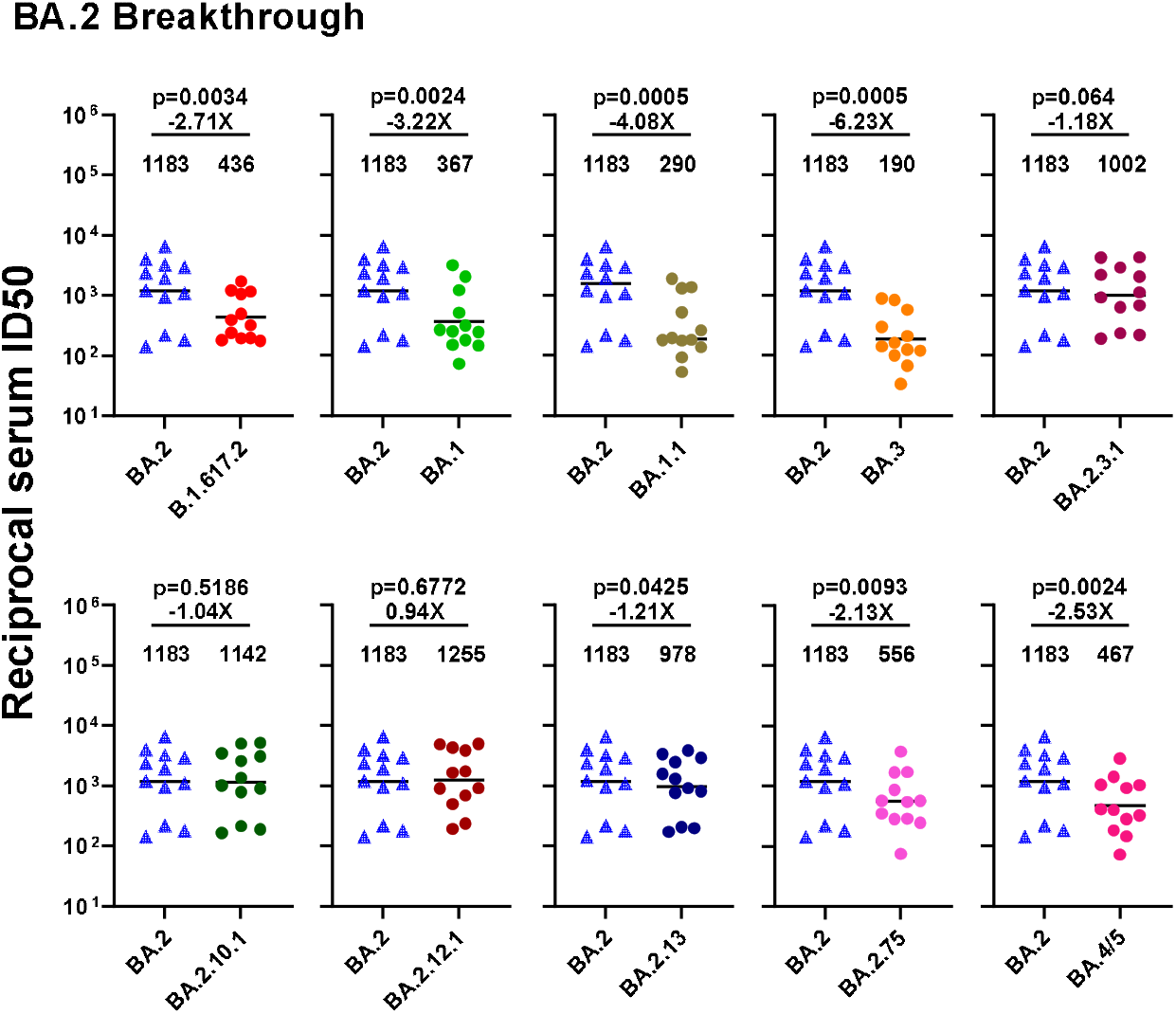
Neutralization activity comparison of BA.2 breakthrough infection sera against BA.2 and other variants.

